# Integrated analysis implicates novel insights of NMB into lactate metabolism and immune response prediction in primary glioblastoma

**DOI:** 10.1101/2025.10.28.685000

**Authors:** Wenjing Bai, Xinglong Li, Yue Li, Peijing Du, Qinghua Zhang, Duo Zheng, Libin Wang

## Abstract

**Background:** Glioblastoma (GBM), the most aggressive primary brain tumor in adults, exhibits profound treatment resistance and poor prognosis. Despite advances in immunotherapy for other cancers, GBM remains refractory, potentially due to its lactate-rich, immunosuppressive tumor microenvironment (TME). While aerobic glycolysis-driven lactate accumulation is known to acidify the TME and impair immune function, the precise mechanisms linking lactate metabolism to immune evasion in GBM remain elusive.

**Methods:** We integrated multi-omics analyses with machine learning to identify - gene signatures associated lactate metabolism (LM) and Tumor Immune Dysfunction and Exclusion (TIDE) through weighted gene co-expression network analysis (WGCNA). Multiple bioinformatic tools were employed to determine the association of key genes with TME remodeling, immunotherapy response and drug sensitivities.

**Results:** Neuromedin B (NMB) emerged as a pivotal regulator of lactate-mediated immunosuppression, correlating strongly with LM and TIDE. We found that NMB expression was remarkably correlated with clinical characteristics, immune response, TME remodeling and drug sensitivities.

**Conclusion:** This study demonstrates that NMB is a key cross-factor in metabolic and immune regulation in GBM. It breaks through the traditional perception that metabolism and immunity are mutually exclusive in cancer by promoting oxidative phosphorylation (OXPHOS) metabolism and enhancing anti-tumor immunity through its dual functions. It provides a new direction for the combined intervention of metabolic targeting and immunotherapy, and the strong predictive value of NMB expression for drug response further highlights its clinical application potential in the formulation of personalized treatment strategies for GBM patients.

## Introduction

Glioblastoma multiforme (GBM), often referred to as grade IV astrocytoma, is the most common and aggressive type of malignant brain cancer in adult^1^. It is characterized by marked invasive, frequent recurrence and poor prognosis posing a major challenge for clinical management^2^. Although surgical resection combined with radiotherapy and chemotherapy can temporarily disease control, the intrinsic drug resistance and immunosuppressive tumor microenvironment (TME) lead to a persistently high failure rate of treatment, with a median survival rate of only12 to 15 months^3-5^. Recent studies have revealed that the immune escape of GBM is closely associated with metabolic reprogramming, particularly aberrant lactic acid metabolism and the dysregulation of immunomodulatory molecules, which jointly shape an immunosuppressive TME and promote tumor progression^6-9^.

The immunosuppressive TME is a marker of glioblastoma (GBM), which contributes to its invasive progression and resistance to treatment^10^. This immunosuppression is primarily mediated by two interrelated mechanisms. Firstly, tumor cells actively secrete immunosuppressive cytokines, including TGF-β and IL-10, which inhibit T cell activation and promote the expansion of regulatory T cells (Tregs)^11, 12^. Secondly, the metabolic reprogramming of GBM cells, especially the Warburg effect (aerobic glycolysis) leads to excessive lactic acid production and accumulation in the TME^13^. The resulting acidic environment polarizes tumor-associated macrophages (TAMs) towards the immunosuppressive M2 phenotype by impairing the functions of cytotoxic T and NK cells, enhancing the immunosuppressive activity of cancer-associated fibroblasts (CAFs), and further strengthening immune evasion^14-17^. In conclusion, these mechanisms establish a self-sustaining immunosuppressive network, promoting tumor immune escape and limiting the efficacy of conventional and immunotherapy interventions. Understanding the interplay between immunosuppressive signaling and metabolic dysregulation is crucial for formulating strategies to overcome GBM immune resistance.

Neuromedin B (NMB), a neuroendocrine peptide acting through the G protein-coupled receptor NMBR (BB1), has emerged as a potential modulator of tumor progression^18, 19^. Emerging evidence indicates that NMB plays a more complex and environment-dependent role in tumor biology. Notebly, NMB exhibits dual functions in TME. It not only promotes the proliferation and invasion of tumor cells via autocrine/paracrine mechanisms, but also modulates inflammatory and immune responses^20, 21^. However, the mechanism involvement of NMB in the immune-metabolic circuitry of GBM, particularly its possible interplay with lactate metabolism, remains largely undefined.

To address this issue, we performed weighted gene co-expression network analysis (WGCNA) on 249 GBM samples and identified NMB as a hub gene co-expressed with immune evasion (TIDE score) and lactate metabolism (LM score) signatures. Comprehensive correlation analyses demonstrated that NMB functions as a pivotal regulator of the metabolism-immune axis in GBM. Mechanistically, NMB promotes oxidative phosphorylation (OXPHOS) and enhances anti-tumor immunity by facilitating T-cell infiltration and function, augmenting antigen presentation and regulating chemokine networks. Furthermore, we have discovered a genetic marker related to lactic acid metabolism, which distinguishes patients based on their metabolic immune profiles, providing a new framework for personalized immunotherapy methods for this refractory disease. Collectively, these findings advance our understanding of the immunosuppressive mechanism of GBM and undercover potential therapeutic targets for overcoming immune resistance.

## Materials and methods

### Data Acquisition and Preprocessing

Transcriptomic data from TCGA-GBM (https://portal.gdc.cancer.gov/) (391 samples: 373 primary tumors, 14 recurrent tumors, 5 normal samples) were obtained via TCGAbiolinks R package. Expression data in TPM format were processed by averaging multi-probe genes. Survival metrics (OS, PFI, DSS) were retrieved from UCSC Xena (https://xenabrowser.net/datapages/). Two CGGA cohorts (http://www.cgga.org.cn/) (693 and 325 samples) were similarly processed with FPKM-to-TPM conversion and log2(TPM+1) transformation. Quality control excluded samples with missing follow-up or survival <30 days. For scRNA-seq analysis, 39 primary GBM samples from 11 GEO datasets (https://www.ncbi.nlm.nih.gov/geo/) meeting inclusion criteria (Homo sapiens, primary/GBM-WT) were curated.

### Systems Biology Analysis of Metabolic-Immune Crosstalk

Tumor immune evasion was quantified using TIDE (http://tide.dfci.harvard.edu/). LM pathway activity was assessed via ssGSEA (GSVA package) using 262 published LM genes. WGCNA (v1.72) was performed on the top 75% variant genes (MAD-filtered) with β=12 soft threshold. Modules were identified (minSize=50, cutHeight=0.25), with eigengenes correlated to TIDE/LM scores. Differential expression analysis (limma, adj.P<0.05, |log2FC|>1) between GBM and normal tissues identified candidate genes intersecting with key modules. KEGG pathway enrichment was performed using clusterProfiler.

### Machine Learning-Based Biomarker Discovery

Univariate Cox regression screened module genes for OS association. LASSO (glmnet) and RSF (randomForestSRC) were applied for feature selection, with consensus genes retained. Hub genes required positive correlation (Pearson r>0, P<0.05) with both TIDE and LM scores.

### Multi-Omics Characterization of NMB

Expression analysis integrated TCGA/GTEx data and HPA IHC images (http://www.proteinatlas.org/). Survival stratification used median NMB expression cutoffs (Kaplan-Meier/log-rank). Genomic alterations were analyzed via cBioPortal (https://www.cbioportal.org/). Methylation profiles were assessed using SMART (http://www.bioinfo-zs.com/smartapp/), evaluating CpG-NMB expression correlations and prognostic value.

### Functional Enrichment

DEGs between NMB^high/low^ groups (|log2FC|>0.585, adj.P<0.05) underwent GO/KEGG analysis (clusterProfiler). Pathway activity was quantified via PROGENy using MSigDB gene sets (GO/KEGG/HALLMARK) (https://www.gsea-msigdb.org/gsea/msigdb/index.jsp). GSEA (fgsea, 1000 permutations) identified enriched pathways.

### Single-Cell and Spatial Transcriptomics

Single-cell RNA sequencing (scRNA-seq) data processing was performed using Seurat (v5.2.0). Quality control filters removed genes detected in <5 cells and cells with <300 expressed genes, <500 total counts, or >20% mitochondrial content. Additional filtering excluded dissociation-related genes (mitochondrial, ribosomal, hemoglobin, and heatshock genes). Data normalization, highly variable gene (HVG) identification, and principal component analysis (PCA) were performed using Seurat (v5.1.0), followed by batch correction with Harmony. Clustering analysis was conducted using the first 20 principal components at a resolution of 0.5, with cell type annotation performed via singleR (v1.2.0) referencing HumanPrimaryCellAtlas, BlueprintEncode, and ImmuneCellExpression datasets, supplemented by manual annotation using canonical markers. Metabolic pathway activity was quantified using the scMetabolism R package with AUCell scoring. Spatial transcriptomics data (GSE194329) were processed similarly and visualized using SpatialFeaturePlot.

### Tumor microenvironment immunophenotyping

Immune landscape characterization included: (1) expression analysis of 122 immunomodulators and 20 immune checkpoint inhibitors from published resources; (2) tumor-infiltrating immune cell (TIIC) estimation using nine algorithms (MCPCOUNTER, EPIC, XCELL, QUANTISEQ, CIBERSORT-ABS, CIBERSORT, TIMER(http://timer.comp-genomics.org/timer/), TISIDB, and TIP (http://biocc.hrbmu.edu.cn/TIP/)); (3) cancer-immunity cycle scoring via the TIP web portal; and (4) single-sample gene set enrichment analysis (ssGSEA) of 28 immune cell types. Correlations between NMB expression and immune features were systematically evaluated.

### Therapeutic response prediction

Immunotherapy sensitivity was assessed using six established gene signatures (Ayers.INFG.SIG, Ayers.T.cell.SIG, Rooney.Cytotoxic.SIG, Zhang.Stem.SIG, Feng.MHC.SIG, and Chen.LM.SIG), with adaptation of the Stem.SIG to include 450 detectable transcripts. Drug sensitivity profiles were predicted using oncoPredict (v1.2) with GDSC2 and CTRP2 datasets, focusing on compounds showing significant correlations (|r| > 0.5, P < 0.05) between ln(IC50/AUC) values and model gene expression.

### Statistical analysis

Associations were evaluated using Spearman/Pearson correlation tests. Group comparisons employed Wilcoxon rank-sum tests. Survival analyses utilized Cox proportional hazards models and Kaplan-Meier methods with log-rank tests. All analyses were performed in R (v4.5.0) with significance thresholds: *p < 0*.*05, p*≤ *0*.*01, p*≤ *0*.*001, p* ≤ *0*.*0001*.

## Results

### 1. WGCNA and Machine learning identifies NMB was associated with TIDE and LM in primary GBM

This study analyzed the transcriptome data of 249 GBM samples using WGCNA, identifying 13 co-expression modules (Figure. 1A-B). Among them, the darktturquoise module was significantly positively correlated with tumor immune dysfunction and rejection (TIDE score: R = 0.81, p = 6e-58) and lactate metabolism (LM score: R = 0.30, p = 2e-06) (Figure. 1B), and was enriched in the oxidative phosphorylation pathway (Figure. 1C). Through a five-step screening strategy (differential expression analysis, module gene intersection, survival correlation analysis, LASSO-Cox/RSF model, and trait correlation verification), TMEM176B, NMB, and RPL39 were finally determined as key genes simultaneously associated with TIDE and LM scores (Figure 1D-H). NMB exhibits a unique protective effect in GBM: patients with high expression of NMB have longer overall survival (HR = 0.676, P = 0.004), disease-free survival (HR = 0.655, P = 0.003), and progression-free survival (HR = 0.722, P = 0.003) (Figure 1I). Further analysis revealed that NMB expression was positively correlated with immune escape (R = 0.353, P < 0.001) and lactate metabolism (R = 0.53, P < 0.001) (Figure 1J-K), suggesting that NMB play a dual role in GBM by regulating immune metabolic reprogramming.

**Figure 1.**
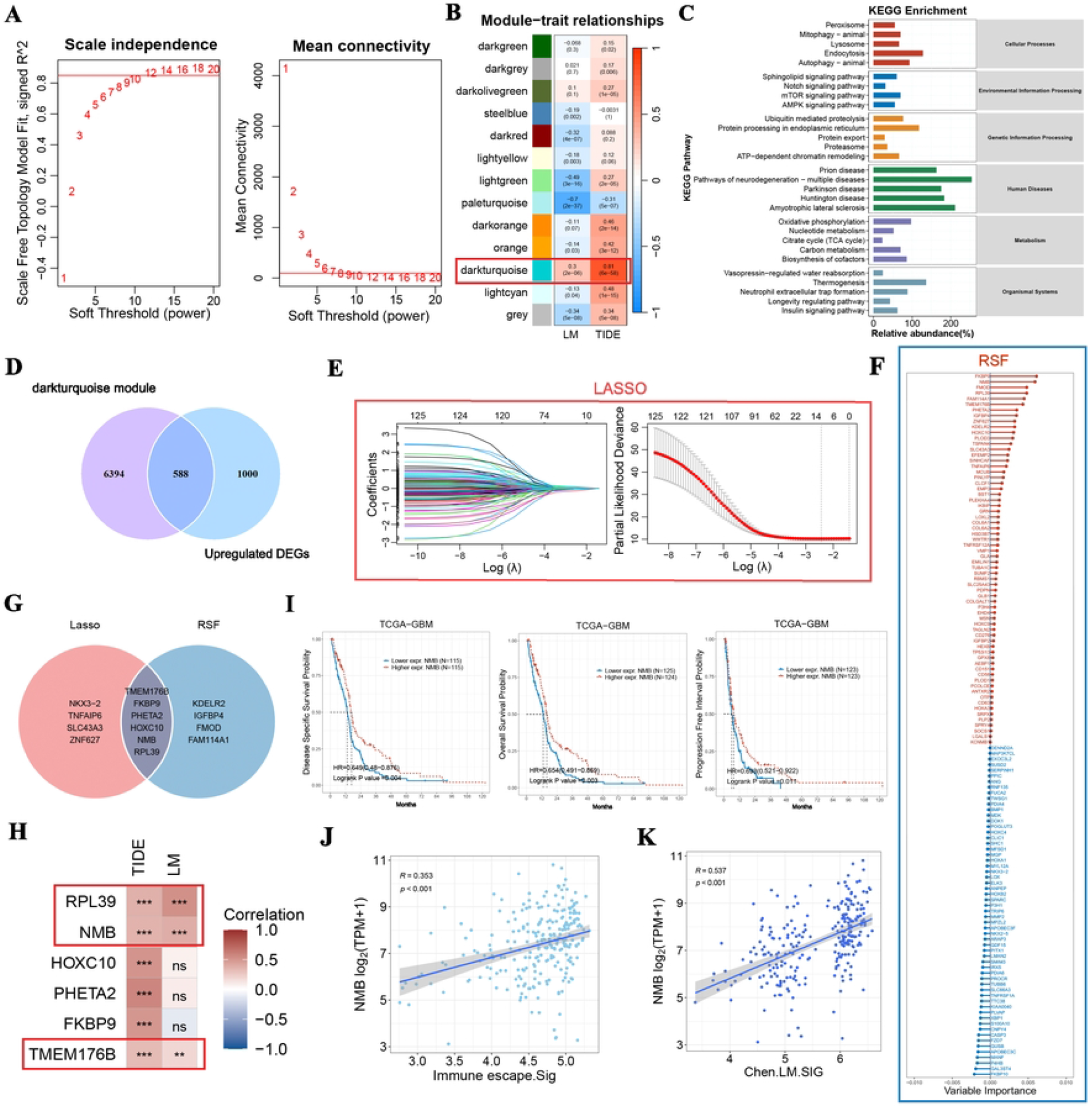
WGCNA and Machine learning identifies NMB was associated with TIDE and LM in primary GBM. (A, B) GBM data from the TCGA database were used to construct gene modules with similar expression patterns to TIDE score and LM score, and cluster analysis categorized into 13 modules. (C) Heatmap shows the correlation of each module with both TIDE score and LM score. (D) A Venn plot of the intersection of DEGs that are abnormally upregulated in primary GBM and genes within the darkturquoise module. (E, F) The LASSO-Cox analysis and the RSF algorithm respectively identified the key prognostic genes that were highly correlated with the patient’s overall survival rate. (G) Hub genes meeting both LASSO and RSF. (H) Heatmap of the correlation between hub genes and TIDE score as well as LM score. (I) Survival curves of OS from two CGGA-GBM cohorts. (J, K). The correlation analysis of NMB expression with immune escape and LM was conducted using the TCGA-primary GBM cohort.

### 2. NMB Displays High Expression and Specific Cellular Localization in GBM

To further validate the expression characteristics of NMB in GBM, this study analyzed the integrated TCGA and GTEx databases, revealing that the expression of NMB in GBM tissues (including different histological subtypes) was significantly higher than that in normal brain tissues (Figure 2A). The immunohistochemical results from the HPA database further confirmed that NMB was mainly located in the cytoplasm and cell membrane, and its expression level in high-grade gliomas was significantly higher than that in the normal cerebral cortex (Figure 2B). To circumvent the spatial resolution constraints of bulk sequencing approaches, we analyzed over 49,000 cells from two single-cell datasets (GSE271379 and GSE256490). We found that NMB was specifically highly expressed in tumor cells and TAMs, but nearly absent in oligodendrocytes, pericytes, endothelial cells, and T/NK cells (Figure 2C-D). These results consistently indicate that NMB shows a significantly upregulated expression pattern in GBM tumor cells and specific immune cell subsets, laying a molecular foundation for its functional study in the GBM microenvironment.

**Figure 2.**
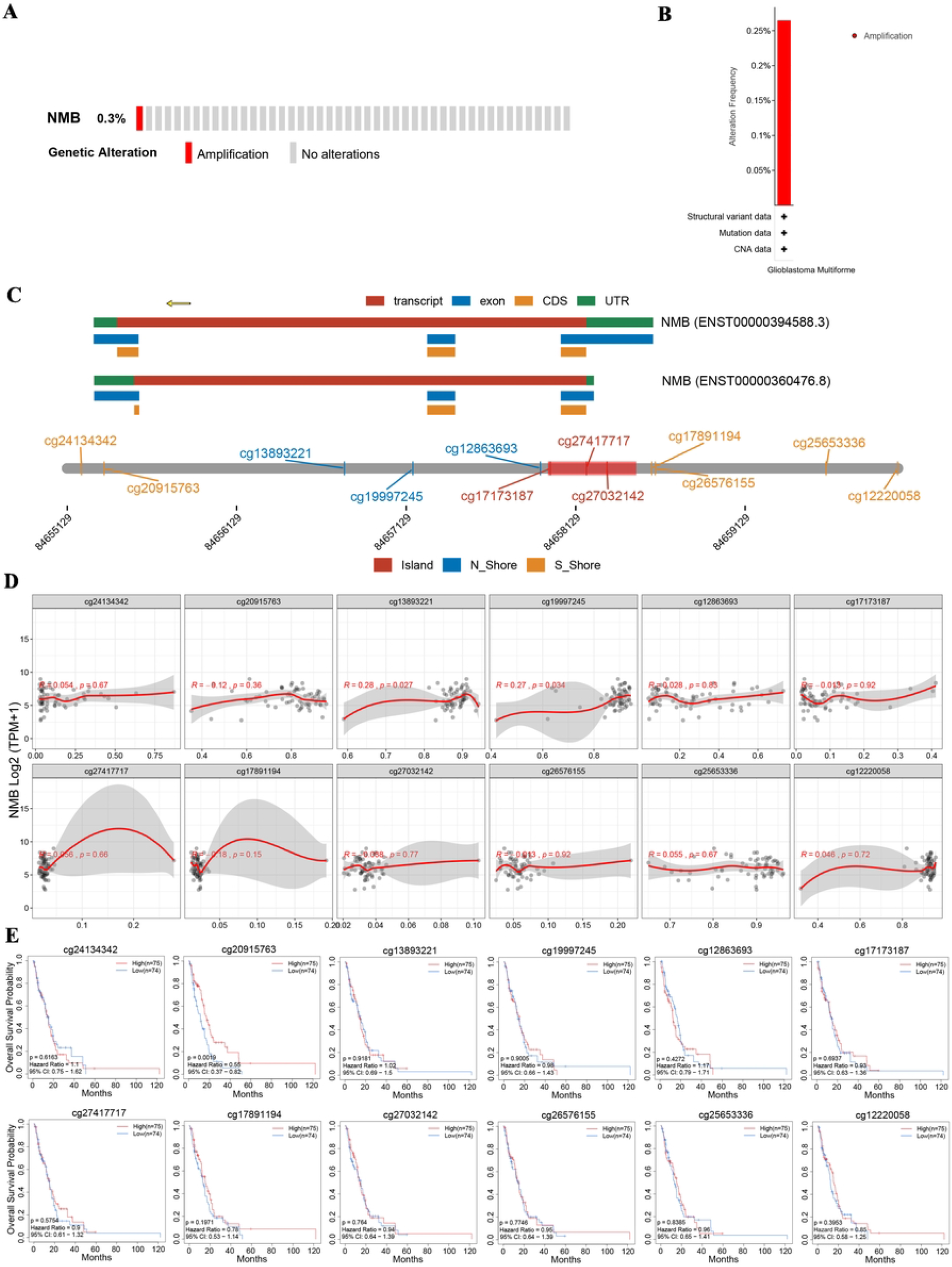
NMB is highly expressed in tumors and TAMs. (A) The expression of NMB between NB and GBM tissues as well as GBM tissues of different histological types. (B) Protein levels of NMB in normal cerebral cortex and GBM samples was visualized by immunohistochemistry via the HPA database. (C-D) Prognostic relationship (OS, DSS, and DFS) between NMB and patients with primary GBM in TCGA-GBM cohort by the univariate Cox regression analysis, Kaplan-Meier analysis, and log-rank test.

### 3. The site mutations of NMB in GBM are closely related to the survival rate of patients

To elucidate the mechanisms underlying NMB overexpression in GBM, we performed comprehensive genomic and epigenomic analyses using TCGA data. Genetic alteration profiling revealed an overall low mutation frequency of NMB (1%) in GBM, with copy number alterations (CNAs) restricted solely to amplification events (Figure 3A, B). However, the observed amplification rate was remarkably low (0.26%), suggesting that CNA-mediated gene dosage effects are unlikely to be the predominant driver of NMB overexpression in GBM.

**Figure 3.**
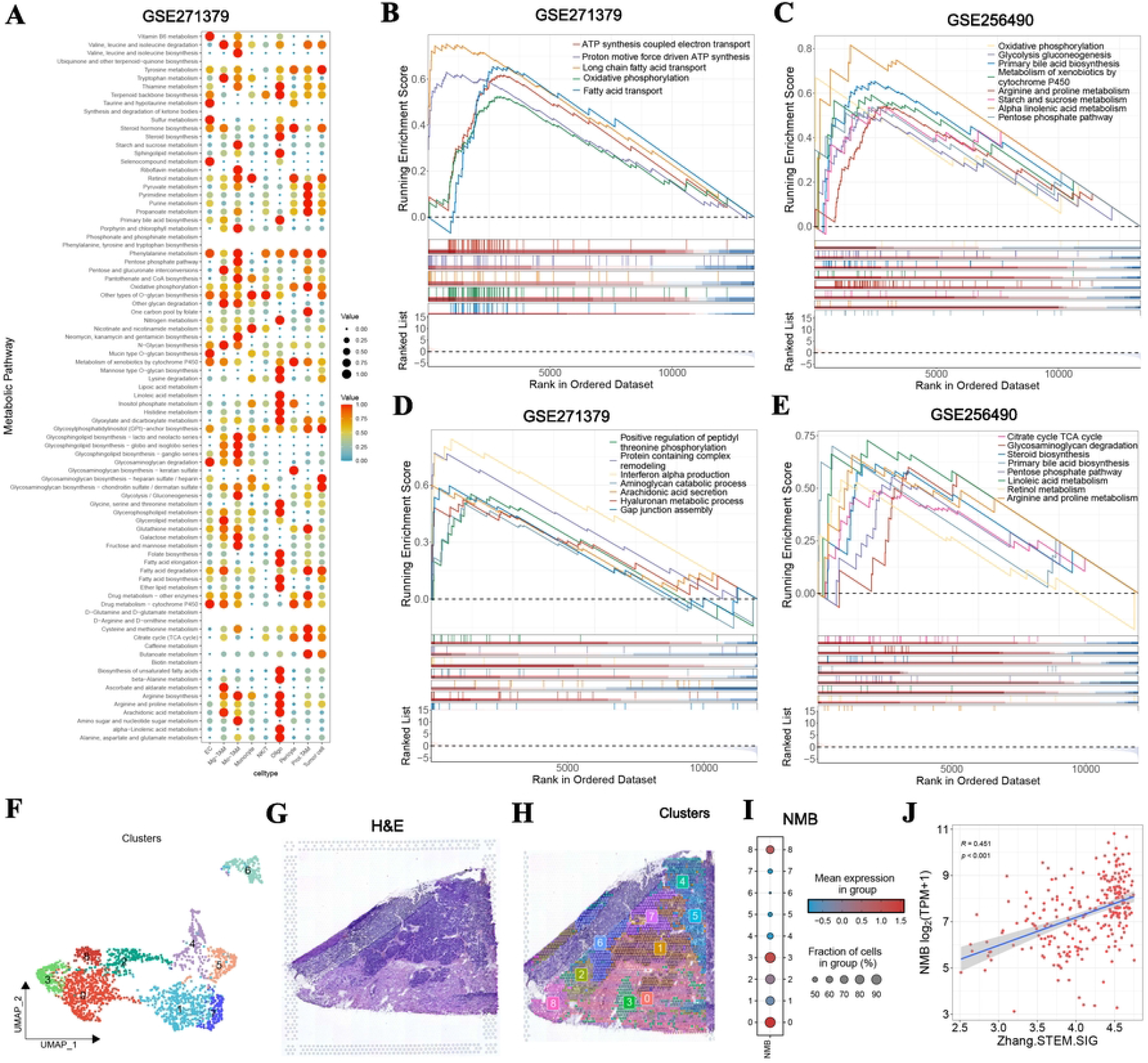
Genetic mutation, and methylation alteration of NMB in GBM. (A-B) The mutation frequency, type and frequency of NMB genomic alterations in GBM obtained from the cBioPortal database. (C-D) Exploration of the detailed chromosomal distribution of methylation probes associated with NMB and the potential association between NMB and the methylation status of CpG islands in GBM were conducted using the SMART App. (E) Prognostic value of NMB-specific CpG site methylation in GBM.

Epigenetic analysis identified 12 CpG sites within the NMB locus, exhibiting an average methylation level (β-value) of 0.419 (Figure S1A, B). While global methylation patterns showed no significant correlation with NMB expression (R=0.1, P=0.43), two specific CpG sites - cg13893221 (R=0.27, P=0.033) and cg19997245 (R=0.26, P=0.036) - demonstrated positive associations with mRNA levels (Figure 3D), implicating their potential role in transcriptional regulation.

Survival analysis revealed a complex relationship between NMB methylation and clinical outcomes. Although overall methylation levels were not prognostic, hypermethylation at cg20915763 was significantly associated with reduced overall survival (Figure 3E). These findings suggest that while NMB overexpression in GBM is not primarily driven by genetic alterations or global methylation changes, site-specific epigenetic modifications may contribute to its dysregulation and influence patient prognosis.

### 4. The high expression areas of NMB have active metabolism and enhanced cell stemness

The metabolic heterogeneity of different cell types in the TME is crucial for elucidating the mechanisms of tumor progression and developing targeted therapies^22^. In the study of GBM, traditional bulk RNA sequencing cannot resolve the metabolic heterogeneity of cells in the TME. Therefore, through single-cell transcriptomic techniques (combined with scMetabolism analysis) and public datasets (GSE271379, GSE256490), we discovered that tumor cells and proliferative macrophages (Prol.TAM) have specific high-activity oxidative phosphorylation (OXPHOS) and tricarboxylic acid cycle metabolic pathways (Figure 4A). In order to analyze the role of NMB in tumor cells, we divided the tumors into the NMB high-expression group and the NMB low-expression group based on the expression level of NMB. The results of the GSEA analysis of the GO-BP item revealed that the tumor cells in the high-expression group were significantly enriched in energy metabolism processes such as electron transfer coupled ATP synthesis, proton-driven ATP synthesis, and long-chain fatty acid transport. (Figure 4B, C). Meanwhile, KEGG analysis showed that metabolic pathways such as OXPHOS, glycolysis/glycogenolysis, and pentose phosphate pathway were significantly activated (Figure 4D, E). Spatial transcriptome localization confirmed that the expression of NMB was significantly increased in the 0, 3, and 8 subgroups of GBM tissues (mainly distributed in the tumor core area) (Figure 4F-I), and was positively correlated with the tumor stem cell characteristics (Figure 4J). These results reveal that the high-expression regions of NMB maintain the characteristics of tumor stem cells by synergistically activating multiple metabolic pathways, and may be an important driving factor for the invasion, metastasis, and drug resistance of the GBM core area.

**Figure 4.**
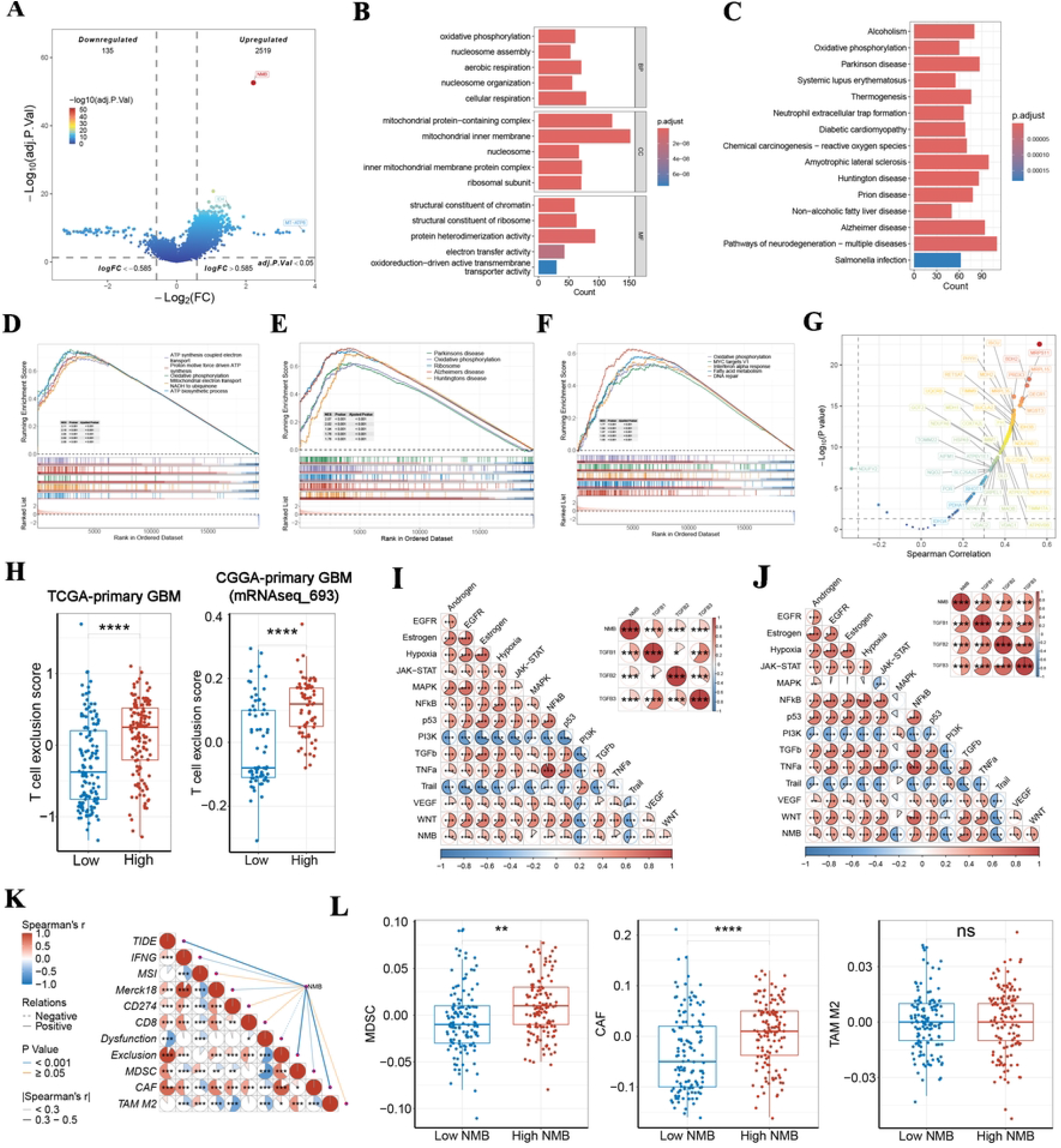
NMB in primary GBM via single-cell sequencing and spatial transcriptome. (A) Bubble plots presenting the metabolic activities of different clusters in GSE271319 cohort. (B, C) GSEA analysis of GO-BP term indicate activities of metabolic related pathways between different NMB groups in tumor cells subtypes. (D, E) GSEA analysis of KEGG term indicate activities of metabolic pathway between different NMB groups in tumor cells subtypes. (F-I) UMAP plot and spatial plots showing the 9 clusters, prediction scores of NMB in tumor region, identified by H&E staining (GEO database, sample ID: GSM5833537, human primary GBM tissue) and original literature’s annotations, as well as the expression intensity of NMB for different clusters. (J) Correlation analysis of NMB expression and Zhang.Stem.SIG.

### 5. NMB is associated with the metabolic reprogramming and formation of immunosuppressive TME in GBM

To further explore the metabolic regulatory role of NMB in GBM and its impact on the tumor microenvironment, the results of this study based on the TCGA and CGGA databases showed that there were 2230 differentially expressed genes (2519 upregulated and 135 downregulated) between the high-expression and low-expression groups of NMB. (Figure 5A). Functional enrichment analysis indicated that NMB-highly-expressed GBM cells were significantly enriched in energy metabolism pathways such as OXPHOS, mitochondrial electron transport chain, and ATP synthesis (Fig. 5B-F), and were positively correlated with core genes of OXPHOS (such as ATP5F1D, NDUFA3, etc.), while negatively correlated with branched-chain amino acid metabolism genes (such as BCKDHA, etc.) (Figure 5G). This suggests that NMB may promote tumor metabolic adaptability by enhancing OXPHOS function and inhibiting branched-chain amino acid metabolism. Additionally, the high-expression group of NMB exhibited significant T-cell exclusion characteristics (Fig. 5H), and was closely related to increased activation of the TGF-β pathway, CAF and myeloid-derived suppressor cells (MDSC) infiltrations (Fig. 5I-L). These findings reveal the dual role of NMB in GBM: it drives mitochondrial metabolic reprogramming to meet the energy requirements of the tumor, and shapes an immunosuppressive microenvironment to promote immune escape, indicating that NMB may be a key immune-metabolic regulatory molecule in GBM.

**Figure 5.**
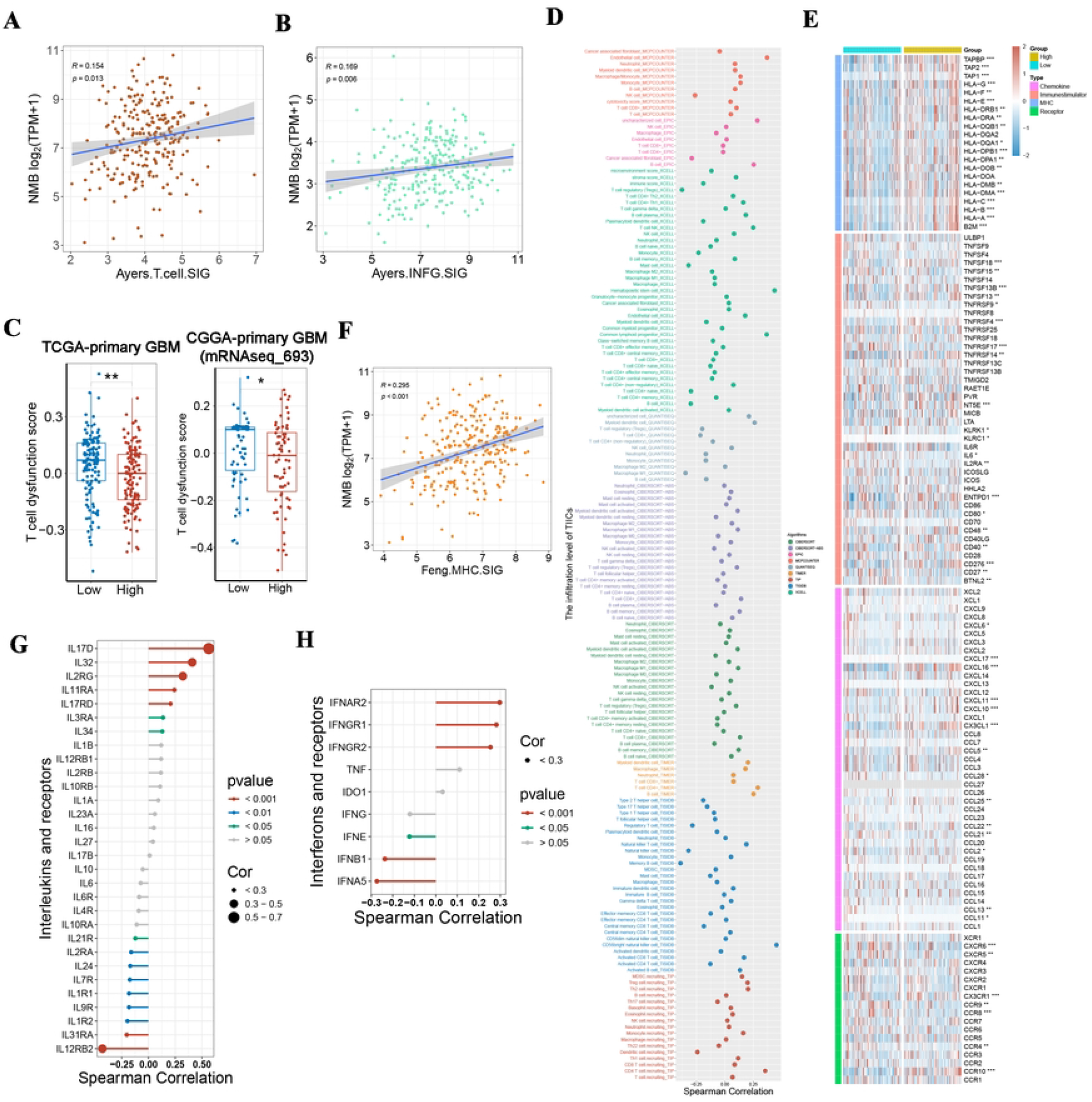
NMB is associated with the metabolic reprogramming and formation of immunosuppressive TME in primary GBM. (A) The volcano plot of 2,230 DEGs. (B) Top 5 enriched terms in the BP, CC and MF based on GO analysis. (C) Bar chart of the top 15 enriched signaling pathways by KEGG enrichment analysis. (D-F) Top 5 enriched terms in the (D) GO-BP, (E) KEGG and (F) HALLMARK by GSEA. (G) Correlation between NMB and 200 markers of OXPHOS. (H) Comparison of T cell exclusion score between the *NMB*^high^ group and *NMB*^low^ group in TCGA-primary GBM and CGGA-primary GBM (mRNAseq_693) cohorts. (I) The potential association between NMB and dysregulated oncogenic pathways as well as TGFB1 (TGF-β1), TGFB2 (TGF-β2), and TGFB3 (TGF-β3) in TCGA-primary GBM cohort and (J) CGGA-mRNAseq_693 primary GBM cohort. (K) Correlation between NMB and immunosuppression scores from TIDE database. (L) The scores between the *NMB*^high^ group and *NMB*^low^ group and immunosuppressation-related cells (MDSC, CAF and TAM-M2).

### 6. NMB is associated with reshaping the TME in primary GBM

The regions with high expression of NMB exhibited higher metabolic activity and characteristics of stem cells (Figure 4). However, patients with high expression of NMB showed a longer survival period (Figure 1I). Therefore, we conducted further in-depth exploration of the mechanism of action of NMB. The study found that the expression level of NMB was significantly positively correlated with the number of tumor-infiltrating T cells and the key markers of T cell function (IFNG) (Figure 6A, B). Further analysis showed that the degree of T cell dysfunction in the high NMB expression group was significantly lower than that in the low expression group (Figure 6C), suggesting that NMB may enhance the anti-tumor activity of T cells. In terms of immune regulatory mechanisms, NMB was positively correlated with antigen-presenting molecules (B2M, HLA-A/B/C, etc.), co-stimulatory molecules (TNFRSF4, CD276), and immune regulatory factors (NT5E, TNFSF13B) (Figure 6D-H), and was significantly associated with the expression of T cell chemokines (CXCL10/11, CCL25, etc.) (Figure 6E). These results indicate that NMB regulates tumor antigen presentation and immune-stimulating signals, drives the infiltration of immune cells, and collaboratively remodels the tumor microenvironment, thereby inhibiting tumor progression and improving patient prognosis.

**Figure 6.**
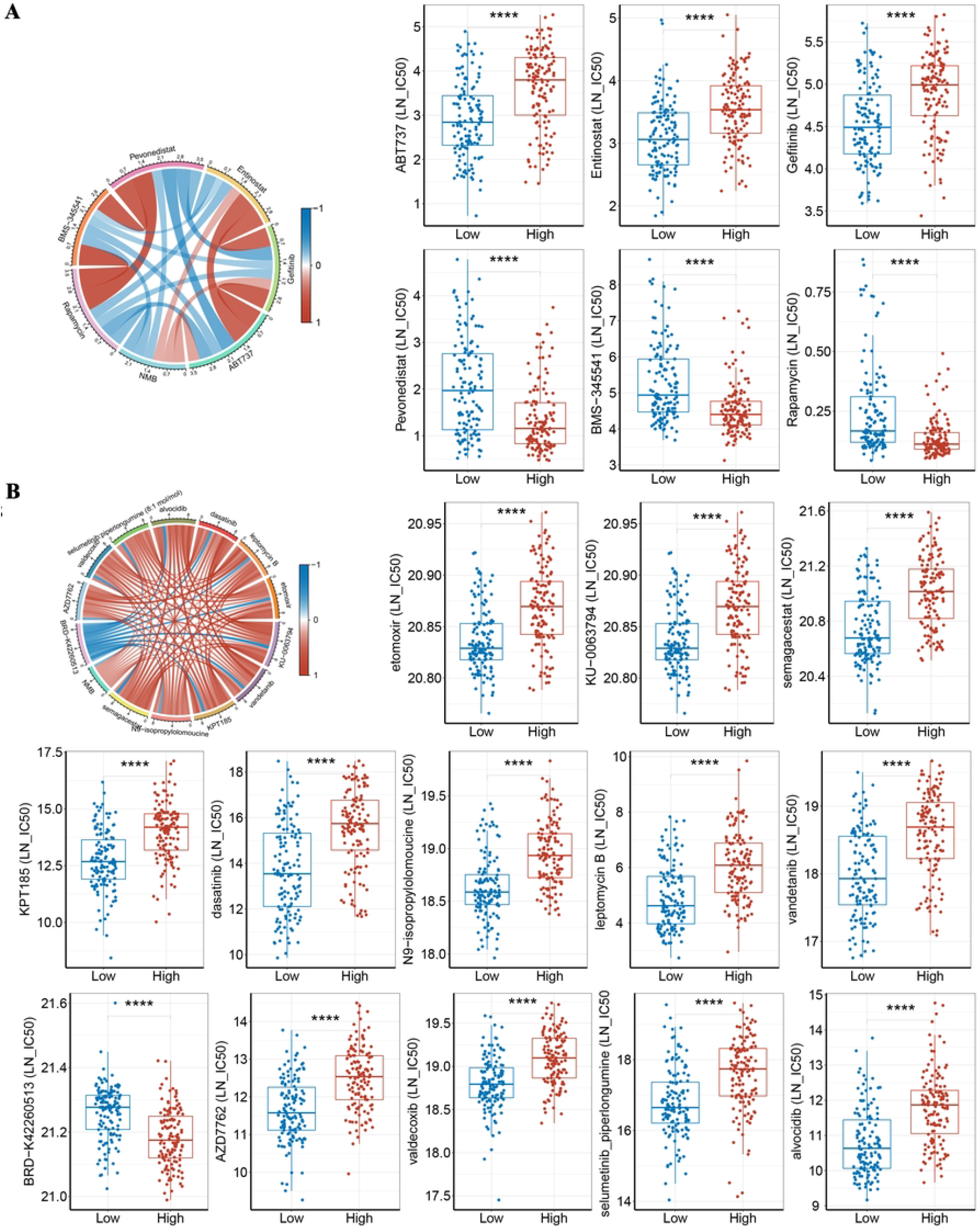
NMB associated with reshaping the TME in primary GBM. (A-B) The association between NMB expression and published predictive signatures related to immunotherapy prediction. (A) Ayers.INFG.SIG, (B) Ayers.T.cell.SIG. (C) Comparison of T cell dysfunction score between the *NMB*^high^ group and *NMB*^low^ group in TCGA-primary GBM and CGGA-primary GBM (mRNAseq_693) cohorts. (D) The relationship between NMB and the infiltration levels of TIICs were calculated using nine independent algorithms. (E) Differences in the expression of immunostimulators, MHC, chemokines, and receptors in *NMB*^high^ and *NMB*^low^ groups. (F) Correlation analysis of NMB expression and Feng.MHC.SIG in TCGA-primary GBM cohort. (G, H) Correlation analysis between NMB expression and (G) interleukins or (H) interferons, as well as their corresponding receptors.

### 7. NMB Serves as a Predictive Biomarker for Anti-Cancer Drug Sensitivity

Based on the XXX role of NMB in GBM, we hypothesized that its expression level can modulate tumor cell sensitivity to anti-cancer agents. Therefore, we systematically analyze the GDSC2 and CTRP2 databases and reveal significant correlations between NMB expression and drug responses. In GDSC2, the expression of NMB was significantly correlated with the LN_IC50 values of six chemotherapy drugs (|R| > 0.4, P < 0.05): patients with low NMB expression were more sensitive to EGFR inhibitor gefitinib, apoptosis inducer ABT737, and histone deacetylase inhibitor entinostat; while patients with high NMB expression showed better therapeutic responses to PI3K/mTOR inhibitor rapamycin, IKK inhibitor BMS-345541, and NEDD8 activating enzyme inhibitor Pevonedistat (Figure 7A). Further analysis of the CTRP2 database revealed that NMB expression levels were significantly correlated with the sensitivity to 13 anti-cancer compounds (|R| > 0.5, P < 0.05): high NMB expression reduced the sensitivity of tumors to γ-secretase inhibitor, CDK inhibitor, nuclear export protein inhibitor, and VEGFR2/EGFR inhibitor, etc., which are broad-spectrum targeted drugs; while patients with low NMB expression showed higher sensitivity to EZH2 inhibitor BRD-K42260513 (Figure 7B). These findings suggest that NMB could serve as a molecular marker to guide individualized chemotherapy in GBM patients.

**Figure 7.**
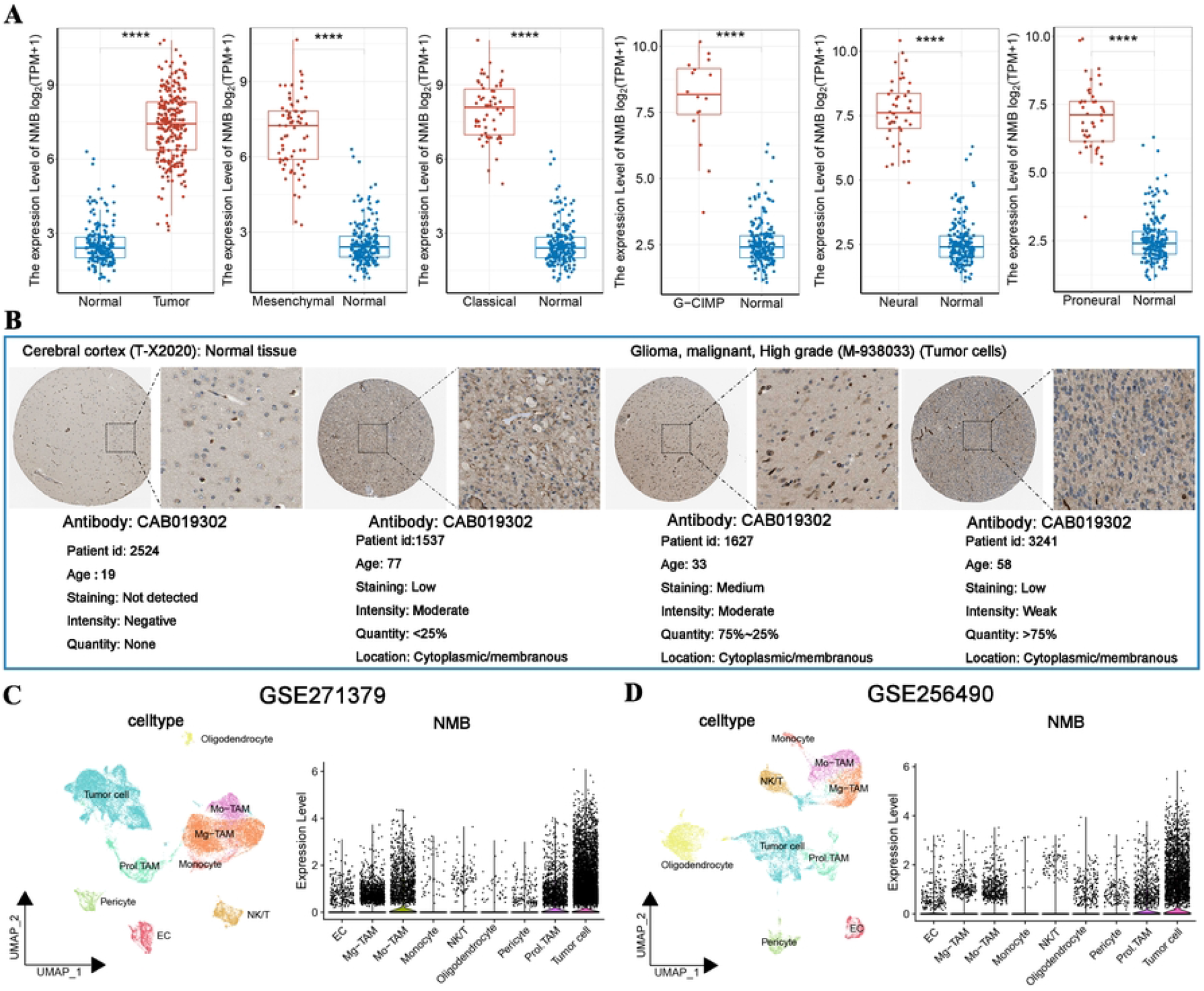
Correlation of NMB expression with the sensitivities of anticancer drugs in primary GBM. (A) Correlation between NMB and the IC50s of seven chemotherapeutic agents in the GDSC2 database, as well as the differences between these drugs across NMB groups. (B) Correlation between NMB and the AUCs of ten compounds in the GDSC2 database, as well as the differences between these compounds across NMB groups.

## Discussion

GBM, the most aggressive primary brain tumor, presents a formidable clinical challenge due to its complex TME characterized by metabolic reprogramming and profound immunosuppression^23, 24^. While the Warburg effect and lactate accumulation have been well-documented in GBM pathogenesis, the molecular regulators connecting metabolic dysregulation with immune evasion remain incompletely understood. Emerging evidence suggests neuropeptides may serve as crucial mediators at this metabolic-immune interface, though their specific roles in GBM remain largely unexplored.

The present study provides a comprehensive characterization of the multifaceted roles of NMB in GBM, revealing its unique involvement in metabolic reprogramming, immune modulation, and therapeutic response. Our findings position NMB as a critical molecular hub that bridges tumor cell-intrinsic metabolic pathways with the immune microenvironment, while also serving as a potential predictive biomarker for targeted therapy in GBM patients.

The most striking finding of our study is the dual nature of NMB’s function in GBM. While NMB acts as a tumor promoter in various other cancers^20^, we observed that high NMB expression in GBM correlates with improved survival outcomes (Figure 1F-G). This paradoxical effect appears to stem from NMB’s ability to simultaneously regulate metabolic reprogramming and immune microenvironment remodeling. Through integrated multi-omics analyses, we demonstrated that NMB-high tumors exhibit a distinct metabolic profile characterized by enhanced OXPHOS and mitochondrial electron transport chain activity (Figure 4A). This metabolic signature differs markedly from the Warburg effect typically observed in cancer cells^23^, suggesting that NMB may drive an alternative metabolic adaptation in GBM that confers both energetic advantages and modulates immune responses.

Our single-cell RNA sequencing analysis provided crucial spatial resolution, revealing that the expression of NMB is particularly abundant in tumor cells and tumor-associated macrophages (TAMs) in the core regions of tumors (Figure 2C, D). This specific localization pattern, combined with the observed positive correlation with tumor stemness markers (Figure 4J), suggests that NMB may play a role in maintaining the stemness characteristics of GBM cells in the core of hypoxic tumors. These observations are further supported by spatial transcriptomics data. Further immune-related analysis revealed that the NMB high-expression group exhibited obvious T-cell exclusion characteristics and was closely related to the increased activation of immunosuppressive cell infiltration in CAF and MDSC (Figure 5H, L). CAF can cause T cells to become marginalized and prevent the process by which T cells gather in the core of the tumor^25^. This is consistent with the phenomenon we analyzed that the high expression of NMB is positively correlated with the content of CAF and the T-cell exclusion is stronger. Furthermore, CAF promotes the secretion of IL-6 through the TGF-β signaling pathway, maintaining the stemness state of the tumor^26, 27^. Most notably, myeloid-derived suppressor cells (MDSC) are a heterogeneous population of immature myeloid cells that can indirectly promote tumor growth and the development of pre-metastatic nics by inhibiting the activity of T cells and NK cells, which is conducive to resistance to immunotherapy^28, 29^. All of these are consistent with the results of the data we analyzed. The above results indicate that NMB plays an important regulatory role in tumor immune escape and metabolism.

The immunomodulatory effects of NMB represent another layer of complexity in its functional repertoire. Contrary to the initial expectations based on its association with immune escape characteristics, high NMB expression is associated with enhanced T-cell chemotactic ability and function, as demonstrated by increased IFNG expression and decreased T-cell dysfunction scores (Figure 6C, E). It is reported that enhancing the antigen presentation ability of tumor cells can not only directly promote T-cell killing but also further strengthen the anti-tumor immune response by releasing more tumor antigens^30, 31^. In addition, the enhancement of immune co-stimulatory signals can amplify the immune response, and the increase in chemokines can also recruit more immune cells to infiltrate the tumor, thereby enhancing the anti-tumor immune effect^32, 33^. Our further analysis revealed that the high expression of NMB can significantly enhance the ability of the antigen presentation mechanism (up-regulating B2M and HLA molecules), while promoting immune stimulation signals (increasing co-stimulatory molecules and chemokines) (Figure 6E-H). These are consistent with the above-mentioned manifestations of enhanced T cell function (Figure 6C). These findings suggest that NMB might create an immune-permissive niche within the otherwise immunosuppressive GBM microenvironment, potentially explaining the improved survival outcomes observed in NMB-high patients.

From a therapeutic perspective, our drug sensitivity analyses yielded clinically relevant insights. The strong correlation between NMB expression levels and response to specific targeted therapies provides a potential framework for treatment stratification. Notably, the differential sensitivity patterns align well with the metabolic characteristics of NMB-high and NMB-low tumors. The preferential response of NMB-high tumors to mTOR and IKK inhibitors corresponds with their OXPHOS-dominant metabolic profile, while the sensitivity of NMB-low tumors to EGFR and HDAC inhibitors matches their likely reliance on alternative metabolic pathways (Figure7A, B). These findings underscore the potential of NMB as a predictive biomarker for personalized therapy in GBM.

Several intriguing questions emerge from our findings. Firstly, the molecular mechanism of the cell type-specific effect of NMB (promoting tumors in other tumors but having a protective effect in GBM) remains unclear. Secondly, the exact pathways by which NMB coordinates metabolism and immune reprogramming deserve further study. Thirdly, the temporal dynamics of NMB during tumor development and treatment response remain unclear. Based on the content of this study, exploring therapeutic strategies targeting the NMB pathway, such as regulating metabolic programming through NMBR, combining it with immune checkpoint inhibitors, and metabolic intervention for OXPHOS with high NMB tumor levels, may become effective means for the treatment of GBM.

In conclusion, our study establishes NMB as a critical regulator at the intersection of metabolism and immunity in GBM. The unique dual functions of NMB in promoting OXPHOS metabolism and enhancing anti-tumor immunity challenge the traditional view that these processes are mutually exclusive in cancer. The strong predictive value of NMB expression for drug response further highlights its potential clinical utility in guiding personalized treatment strategies for GBM patients.

## DATA AND CODE AVAILABILITY

All data have been presented in the manuscript.

## SUPPLEMENTAL INFORMATION

Supplemental information can be found online at https

## AUTHOR CONTRIBUTIONSR

Wenjing Bai and Xinglong Li performed and analyzed the experiments. Wenjing Bai and Xinglong Li designed the study. Wenjing Bai, Yue Li and Peijing Du helped to collect data. Wenjing Bai and Xinglong Li analyzed and interpreted data. Libin Wang and Xinglong Li secured funding. Duo Zheng and Qinghua Zhang supervised the project. Wenjing Bai, Xinglong Li, Duo Zheng, Qinghua Zhang and Libin Wang drafted the manuscript. All authors edited and approved the final manuscript.

## CLINICAL TRIAL NUMBER

Not application.

